# The capability of plant-bacteria consortia to reduce the genotoxicity of unsymmetrical dimethylhydrazine in the environment

**DOI:** 10.1101/2025.01.17.633569

**Authors:** Uliana S. Novoyatlova, Andrew G. Kessenikh, Neonila V. Kononenko, Ekaterina N. Baranova, Stanislav Ph. Chalkin, Sergey V. Bazhenov, Ilya V. Manukhov

## Abstract

Unsymmetrical dimethylhydrazine (UDMH) despite its proven high toxicity continues to be used in rocket technology and some other areas of human activity. In this work, the ability of plant-bacteria consortia to reduce the genotoxicity of UDMH incomplete oxidation products was investigated. Genotoxicity was assessed using a specific lux-biosensor *Escherichia coli* MG1655 pAlkA-lux sensitive to DNA alkylation in cell. For microbiological biodegradation, the *Bacillus subtilis* KK1112 strain was obtained by the isolation from soil with a subsequent selection for resistance to high UDMH concentrations (more than 5000 MAC). Its ability to biodegrade UDMH was shown by observing the reduction of DNA alkylation of the KK1112-treated UDMH. The ability of KK1112 cells to act in a bacterial-plant consortium with following fodder halophytes was studied: *Bromus inermis* Leyss, *Medicago varia* Mart. and *Phleum pratense* L. A synergistic reduction in the alkylating properties of UDMH oxidation products was observed under the combined use of bacteria and plant seedlings. The greatest effect was obtained when bacteria was used in combination with *B. inermis*. It was shown that KK1112 cells accelerate the seedlings development and mitigate the growth inhibition that occurs during incubation with UDMH. The obtained results indicate that it is optimal to introduce the bacterium KK1112 in combination with *B. inermis* plants for soils in arid climate zones during reclamation after UDMH exposure.

## 1. Introduction

UDMH remains the primary contributor to the negative environmental impacts of spacecraft launches [1]. UDMH is a strong reducing agent and can be oxidized by both strong nitric acid-based oxidants (nitrous oxide) and atmospheric oxygen to form several incomplete oxidation products: dimethylamine, tetramethyltetrazene, nitrosodimethylamine (NDMA), methylene dimethylhydrazine, formaldehyde, hydrocyanic acid, nitrogen oxides, and others [2]. The greatest danger of UDMH for living organisms is the genotoxic effects of its incomplete oxidation products, which can appear both inside living cells and when interacting with atmospheric oxygen [3,4]. The most dangerous of these are compounds of the nitrosodimethylamine type, which alkylate the nitrogenous bases in DNA [5].

Many works are devoted to the study of UDMH destruction processes in soils [6–10]. Studies on soil detoxification were conducted using shungite sorbents [11], silicon-containing fertilizers, and lignohumic components [12]. The physicochemical methods for water purification based on complete UDMH oxidation were proposed [13,14]. It was noted that the most intensive processes of UDMH destruction in the soil occur in the zones of plant growth [15].

There are several studies devoted to the microbiological biodegradation of UDMH. Chugunov *et al*. demonstrated the ability of some strains of bacteria, micromycetes, and yeast to biodegrade UDMH [16]. The strains were isolated from tundra soils containated with rocket fuel components, and the work presents data indicating the ability of these microorganisms to utilize UDMH as the only source of nitrogen, carbon, and energy. In the work of American researchers, the possibility of industrial biotreatment of wastewater containing methylhydrazine and hydrazine by bacteria primarily of two genera, *Achromobacter* and *Rhodococcus*, was demonstrated [17]. Representatives of another two genera: *Comamonas* sp. and *Stenotrophomonas* sp., which are capable of biodegrading UDMH under certain conditions were isolated from the activated sludge of the Beijing Qinghe Wastewater Treatment Plant (Beijing, China) [7].

In this work, we investigated the ability of bacteria and plants to reduce the alkylating capacity of UDMH-polluted environment. To assess the DNA-alkylating activity, we used *E. coli* MG1655 pAlkA-lux biosensor cells [3]. The pAlkA-lux plasmid contains the *luxCDABE* genes under the control of the P_*alkA*_ DNA glycosylase promoter, which is activated by the Ada regulator [18]. The *E. coli* pAlkA-lux biosensor allows one to assess the presence and concentration of alkylating substances in the medium based on the level of bioluminescence: the luminescence of cells increases with the increase of the DNA alkylation effect [19,20].

As a result of the search for microorganisms capable of biodegrading UDMH, a UDMH-resistant strain *B. subtilis* KK1112 was isolated from the UDMH-treated soil after long-term selection. The efficiency of this strain used in consortia of plants and bacteria to reduce the genotoxicity of UDMH incomplete oxidation products was assessed.

## 2. Materials and Methods

### 2.1. Strains and plasmids

The *E. coli* K12 MG1655 F-*ilvG rfb*-50 *rph*-1 strain (obtained from VKPM) was used as a recipient strain to obtain biosensor cells by transforming it with the pAlkA-lux and pColD-lux plasmids.

The *B. subtilis* KK1112 strain is a natural isolate obtained in this work by multiple selection for UDMH resistance; it was used in the plant-bacteria consortium.

*B. subtilis* 168 trpC2 (obtained from VKPM) was used as a control for UDMH-resistance experiments.

The pAlkA-lux plasmid [3] is a derivative of the pDEW201 promoter-probe vector [21] and contains a transcriptional fusion of the *E. coli* P_*alkA*_ promoter with the *luxCDABE* reporter genes from *P. luminescens*.

The pColD-lux plasmid [22] is a derivative of the pDEW201, which contains a transcriptional fusion of the P_*cda*_ promoter with the *luxCDABE* reporter genes from *P. luminescens*.

### 2.2. Media and growth conditions for bacteria

Lysogeny Broth (LB) medium (pH 7.5) was used for cultivation of *E. coli* and *B. subtilis* cells. To obtain a solid medium, bacteriological agar was added to a concentration of 1.5%. Ampicillin 100 mg/L was added to the medium for cultivation of *E. coli* MG1655 pAlkA-lux and *E. coli* MG1655 pColD-lux.

*E. coli* cells were grown in test tubes with LB medium in a shaker with constant 200 rpm aeration or on the surface of LB solid medium in Petri dishes at 37°C.

M9 medium for experiments with UDMH was prepared as in [23].

### 2.3. Selection of soil bacteria for resistance to UDMH

Bacteria for further selection for UDMH resistance were collected from three soil samples: one from the pond bank (55.6165807N 37.6133604E), and two samples of urban soil cover, unfertilized and fertilized with imported chernozem (55.6394050N 37.6185800E). Soil samples were added to phosphate buffered saline (Na_2_HPO_4_ - 8.1 mM, KH2PO4 - 1.5 mM, NaCl – 137 mM, KCl - 2.7 mM) in a ratio of 1/10 w/w, containing UDMH 10 g/L. The samples were incubated in 50 mL tubes with hermetically sealed lids, with slow shaking at 60 rpm at room temperature. Microaerobic conditions were provided by atmospheric oxygen, which was initially present in the tubes (the air volume in the containers was 2/3 of the total). After 15 days of incubation, 20 g/L UDMH was added to the samples (the total introduced UDMH was 30 g/L) and the samples were incubated for another 15 days. These probes were plated on LB agar. Several selected clones were then subjected to an additional round of selection on a liquid medium consisting of phosphate-buffered saline and 1% UDMH, which was assumed to be a source of nitrogen and carbon.

### 2.4. Plants and seedling production conditions

The objects of the study were forage grasses: smooth brome (*Bromus inermis* Leyss), hybrid alfalfa (*Medicago varia* Mart.) and timothy grass (*Phleum pratense* L.). The plant seeds were germinated for 4 days in Petri dishes on wet filter paper. For the experiments samples were prepared with UDMH, plants and *B. subtilis* cells in various combinations. When preparing the samples, one seedling of the selected plant and *B. subtilis* KK1112 bacteria (final titer in the sample 5×10^6^ CFU/mL) were added to 5 mL of 0.5% UDMH solution in a glass Petri dish (approximately, diameter was 4 cm) with a lid and incubated with lighting at room temperature for a week. After incubation, the solution was collected and tested for the content of alkylating substances. Also, the titer of live bacteria in the solution was measured by plating on the solid LB medium.

### 2.5. Bioluminescence measurement

180 µl of the fresh *E. coli* MG1655 pAlkA-lux biosensor cell culture was placed into the wells of a 96-well plate or separate 1.5 mL tubes. 20 μL of the studied samples in different dilutions were added to the aliquots of cell culture. Before the addition, studied samples (the liquid obtained after incubation of *B. subtilis* cells and/or seedlings in UDMH solution) were clarified by centrifugation for 5 min at 10,000 g and sterilized by filtration through a 0.22 μm filter. Biosensor cells with added samples were incubated at room temperature for 3-5 hours with periodic measurements of bioluminescence. The total bioluminescence in each well was measured using the SynergyHT multimodal plate-reader (Biotek Instruments, USA). Luminescence of biosensor cells in capeless tubes was measured using Biotox-7BM (BioPhysTech, Moscow, Russia). Luminescence values were expressed in relative light units (RLU), specific for each luminometer. All experiments were performed in 3-5 replicates. The kinetic curves on the graphs representing the results of individual experiments, the results of which were consistent with the others.

## 3. Results

### 3.1. Obtaining a strain resistant to UDMH

The search for microflora capable of biodegrading UDMH and its toxic partially oxidized derivatives was conducted. The first step involved selection of soil microorganisms based on resistance to high (more than 5000 MAC) concentrations of UDMH. For this purpose, soil samples of various types were collected (chernozem, a sample from the shore of a pond, and a sample of urban unfertilized soil cover). The soils were added (1/10 w/w) to a medium consisting of phosphate-buffered saline (without glucose and any nitrogen source) supplemented with 10 g/L UDMH. After 2 weeks of incubation, its concentration was increased by addition of another 20 g/L of UDMH. After incubation for another two weeks, the living bacteria remaining in the samples were plated on LB agar (Figure S1). Several clones isolated in this manner were subjected to an additional round of selection in phosphate-buffered saline and 1% UDMH (without soil) for 15 days, after which plating was carried out.

One of the clones isolated in the result of selection was analyzed in details. The 16S rRNA gene sequence of the KK1112 isolated clone completely coincides with that of *B. subtilis* (DSM 10). Thus, the isolated strain was designated as *B. subtilis*. Figure 1 shows the results of comparison of KK1112 with the laboratory strain *B. subtilis* 168 for resistance to UDMH. For this experiment, *B. subtilis* cells were incubated in liquid M9 medium (without glucose) with 1% UDMH and titer of living cells was daily determined by plating on LB agar.

**Figure 1.**
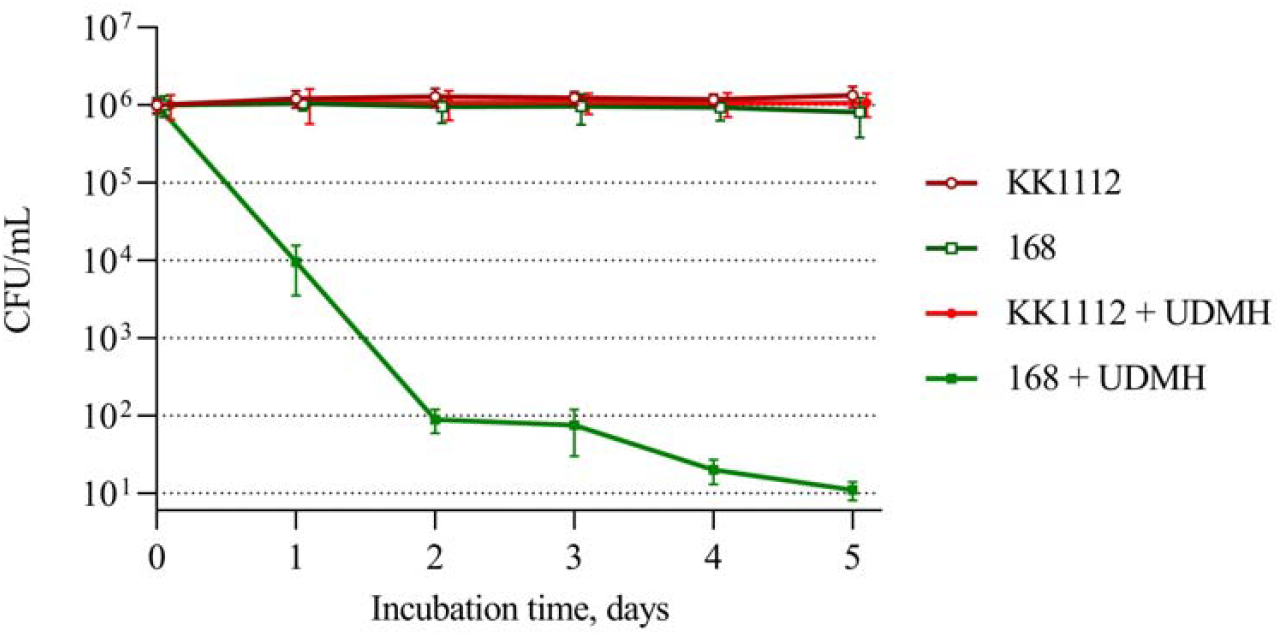
Comparison of *B. subtilis* laboratory strain 168 and selected strain KK1112 for resistance to 0.5% UDMH. Living cell titers are given for samples of cell cultures (“168” and “KK1112”) incubated in medium with 0.5% UDMH (“+UDMH” curves) or without it.

As can be seen from Figure 1, the titer of *B. subtilis* KK1112 does not decrease during incubation with UDMH for 5 days, while the titer of the laboratory strain *B. subtilis* 168 drops to almost zero during this time.

### 3.2. B. subtilis KK1112 cells reduce the alkylating potential of UDMH-containing medium

Figure 3 shows the data on the mutagenicity potential of a medium containing 1% UDMH after weeklong incubation with or without *B. subtilis* KK1112 cells. For this experiment, an overnight culture of *B. subtilis* KK1112 was prepared, pelleted by centrifugation, suspended in M9 medium to a final titer of approximately ∼10^9^ cells/mL, and divided into samples with and without following addition of UDMH. After weeklong incubation, the spent culture medium (SCM) was collected and processed as described in the methods by centrifugation and filter-sterilization, and the resulting sample solutions were examined for genotoxicity using the *E. coli* pAlkA-lux biosensor (Figure 2).

**Figure 2.**
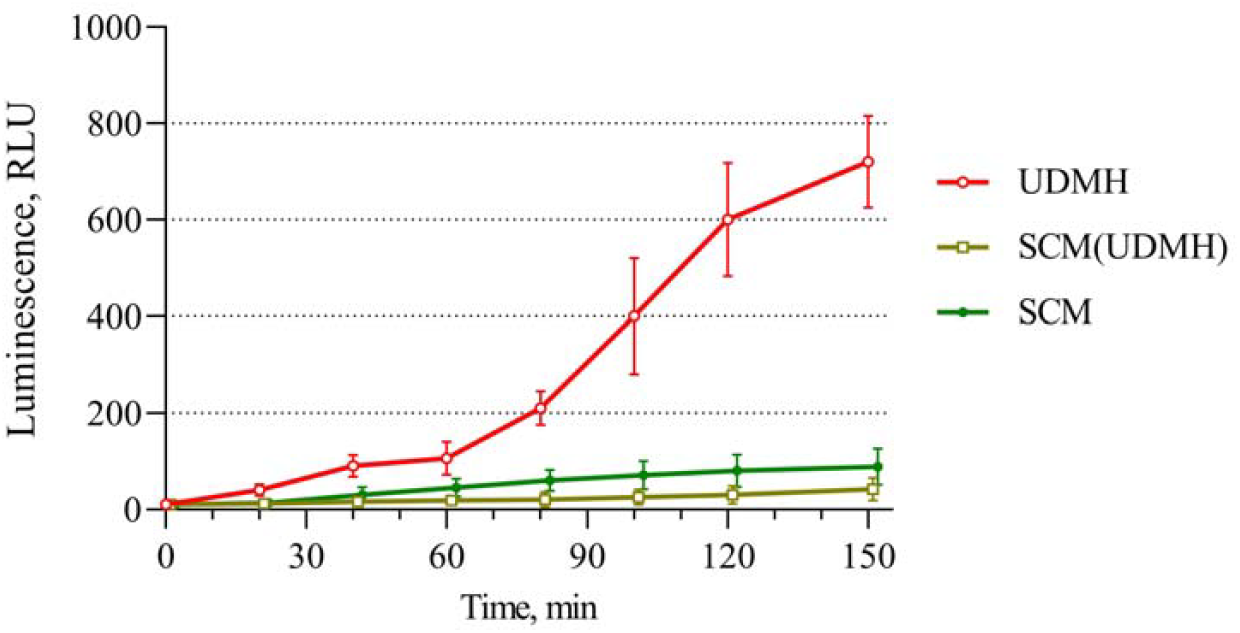
Time dependence of the luminescence of the *E. coli* MG1655 pAlkA-lux biosensor cells in the presence of 1% UDMH pre-incubated for a week in the M9 medium with and without *B. subtilis* KK1112 cells. UDMH – the M9 medium with 1% UDMH after weeklong incubation; SCM – SCM after incubation of *B. subtilis* KK1112 in M9; SCM+UDMH – SCM after incubation KK1112 cells in M9 with UDMH.

**Figure 3.**
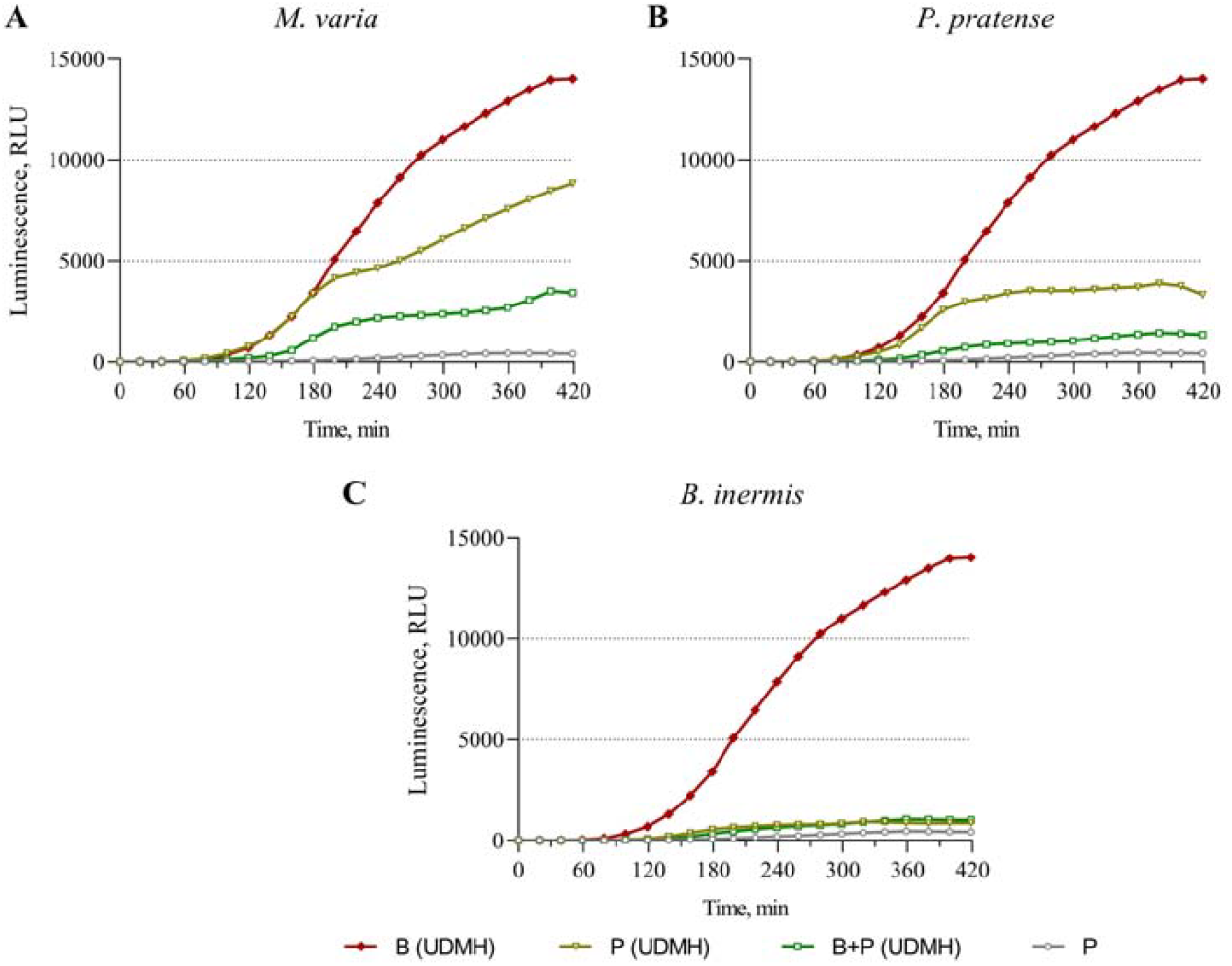
Bioremediation of the UDMH-polluted medium by the plant seedlings and bacteria. Panels A, B, and C correspond to the *M. varia, P. pretense*, and *B. inermis* seedlings, respectively. The graphs show the response of the *E. coli* pAlkA-lux luminescent biosensor cells after the addition of the water solutions supplemented with 0.5% UDMH and incubated during a week in presence of plant seedling (“P (UDMH)” curves), bacterial cells of UDMH-resistant KK1112 strain (“B (UDMH)” curves), and their combination (“B+P (UDMH)” curves). The “P” curve corresponds to a negative control ― luminescence of the *E. coli* pAlkA-lux cells upon addition of the UDMH-free water, in which the same plant seedlings were incubated.

In this experiment, during incubation for a week, UDMH undergoes incomplete oxidation by atmospheric oxygen, forming alkylating agents such as NDMA. An increase in the bioluminescence of the *E. coli* MG1655 pAlkA-lux biosensor cells upon addition of bacteria-free UDMH-containing medium demonstrates the presence of alkylating agents. Addition of *B. subtilis* KK1112 cells (curve “SCM(UDMH)”) to the UDMH-supplemented medium leads to an almost complete absence of the DNA-damaging activity of UDMH oxidation products after a week of incubation.

### 3.3. Synergistic degradation of UDMH by a consortium of bacteria and plant seedlings

To test the ability of plants in consortia with KK1112 bacteria to reduce alkylating activity in a UDMH-polluted environment the following fodder halophytes were selected: smooth brome (*B. inermis* Leyss), hybrid alfalfa (*M. varia* Mart.), and timothy-grass (*P. pratense* L.). The halophytes were chosen in perspective for their practical use in the typical arid zones of spacecraft launches (Baikonur, Kapustin Yar, Semipalatinsk, and others). Their four-day-old seedlings were used in the experiment. To create an effective consortium, *B. subtilis* KK1112 cell culture with a titer of approximately 5×10^6^ was used. Bacteria and plant seedlings were incubated separately and together for a week in Petri dishes with a 0.5% UDMH solution. The resulting solutions were centrifuged, sterilized by filtering, and analyzed for the alkylating capacity using biosensor cells. UDMH-free water in which the same seedlings were incubated during the experiment was used as negative control for subsequent biosensor measurements. The results of the studies of the alkylating capacity of the obtained samples are shown in Figure 3. Three replicates were set for each sample; the figures show the typical curves.

Monitoring the titer of *B. subtilis* KK1112 in samples with UDMH showed that the number of viable bacteria did not decrease during the incubation (Table S1), which was expected based on the previously obtained data on their resistance to UDMH.

The concentrations of UDMH and bacterial titers for these experiments were chosen to ensure that the bacteria could not degrade the alkylating agents during the incubation period without the presence of plants. This is confirmed by the activation of the biosensor when examining a sample obtained by incubating a UDMH solution with KK1112 cells only (curve “B (UDMH)”). One can see that induction of the P_*alkA*_ promoter is enhanced significantly, which results in more than 100-fold rise in the luminescence compared to the control without UDMH, curves “B (UDMH)” and “P” respectively. All plant seedlings demonstrated the ability to bioremediate UDMH-supplemented solution, which is especially noticeable if KK1112 cells were used in low titer. What’s more important is that the combined effect of bacteria and seedlings on reducing alkylating activity turned out to be higher than separately for all kinds of plants in the experiment.

Alfalfa seedlings quite effectively reduce the alkylating properties of the UDMH-polluted environment (Figure 3A); the “P (UDMH)” curve is approximately 1.5 times lower than the “B” curve, which corresponds to the addition of *B. subtilis* KK1112 solely. The combined action of bacteria and alfalfa seedlings decreases the alkylating potency of the medium better, but we can still observe an activation of the biosensor. The effect of the timothy grass seedlings turned out to be stronger in comparison with alfalfa (Figure 3B). It can be estimated by the decrease in the activation of the *E. coli* MG1655 pAlkA-lux biosensor luminescence: the curve “P (UDMH)” in Figure 3B is approximately half as low as in Figure 3A. Solution obtained after incubation of timothy seedlings together with bacteria in UDMH-containing water only slightly activate the biosensor (curve “B + P (UDMH)”). Smooth brome seedlings without the addition of bacteria turned out to be a very effective destructor of UDMH: the alkylating potency of the sample after the incubation decreased almost to the biosensor’s lower limit of detection even without the addition of *B. subtilis* KK1112 (Figure 3C). Addition of *B. subtilis* KK1112 cells to the seedlings enhances this effect even more. It could be seen from the middle part of curve “B + P (UDMH)”, which is lower than curve “P (UDMH)” at a time near 180 min.

### 3.4. Effect of B. subtilis KK1112 cells and UDMH on seedling development

In the conducted experiments, suppression of seedling formation was noted when UDMH was added to the incubation medium. Partial or complete restoration of seed growth and germination was noted in samples containing *B. subtilis* KK1112. Figure 4 shows photographs of seedlings taken after the experiment with the addition of UDMH and *B. subtilis* KK1112 cells, as well as the results of seedling length measurements.

**Figure 4.**
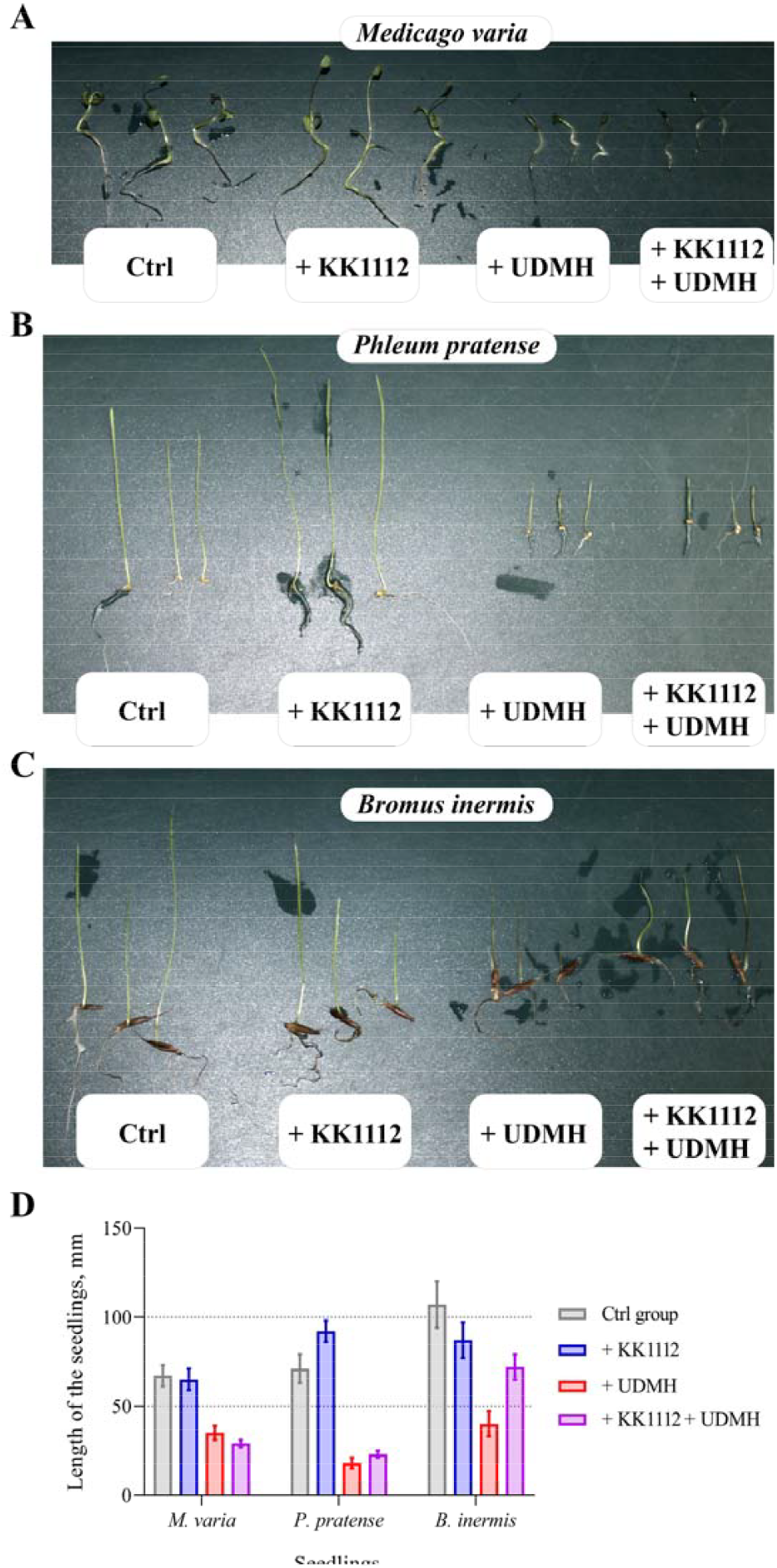
Photos of seedlings of hybrid alfalfa *M. varia* M. (**A**), Timothy-grass *P. pratense* L. (**B**), and smooth brome *B. inermis* Leyss (**C**); and results of seedlings length measurements (**D**). Seedlings were obtained in a result of a weeklong incubation in the pure saline solution (Ctrl) or saline solution supplemented with *B. subtilis* KK1112 cells (+ KK1112), UDMH 0.5% (+ UDMH), or a combination of UDMH and *B. subtilis* KK1112 cells (+ KK1112 + UDMH).

The effect of UDMH on the seedlings of all the studied plants is clearly visible both in the presence of *B. subtilis* KK1112 cells and without them. In all cases, growth inhibition and a change in the shape of the seedlings were observed. It is important to note that this effect is maintained even for brome seedlings, even though they almost eliminate the alkylating effect of the UDMH-containing medium (Figure 3).

The addition of *B. subtilis* KK1112 cells has a different effect on the development of seedlings. The consortium of *M. sativa* seedlings with *B. subtilis* KK1112 bacteria turned out to be the least effective in our experiments. Seedling development was inhibited by UDMH and was practically not restored by adding cells. No positive effect of *B. subtilis* KK1112 on the development of *M. varia* seedlings in the control was also noted. The KK1112 cells had a significant positive effect on the development of timothy-grass *P. pratense* seedlings in the absence of UDMH. The slight positive effect of *B. subtilis* KK1112 cells on the development of *P. pratense* seedlings remains even if UDMH is added to the medium. Although the seedlings became only slightly taller, their better growth can be judged by the thickening of the stem. The best results were obtained for the smooth brome: the addition of *B. subtilis* KK1112 cells nearly eliminates the inhibitory effects of UDMH on the development of brome seedlings. This was evident from both the elongation and thickening of the *B. inermis* stems.

It is known that the UDMH incomplete oxidation products can induce the SOS response in bacterial cells [4]. Therefore, to further test the synergistic effect of the bacteria with brome seedlings, we used the *E. coli* pColD-lux biosensor. This biosensor is sensitive to DNA damage that stalls the replication fork and induces the SOS response [22,24]. Figure 5 shows the luminescence kinetic curves for *E. coli* pColD-lux biosensor cells, the measurements were performed under the same conditions and with the addition of the same samples, as for Figure 3.

**Figure 5.**
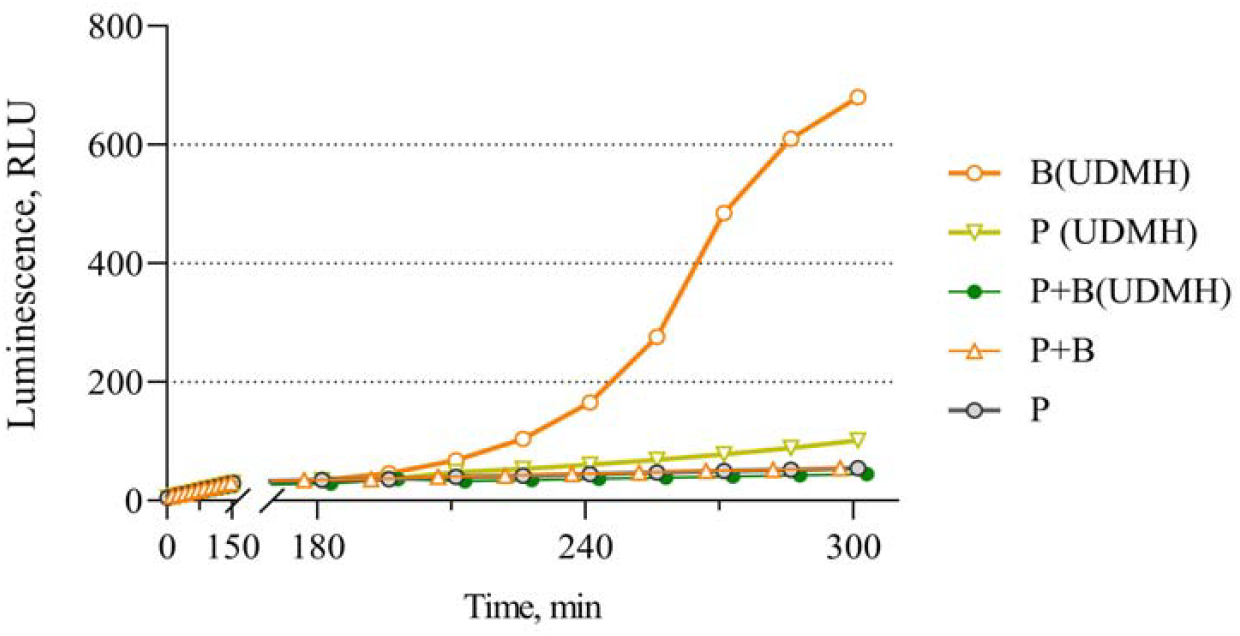
Effect of *B. inermis* seedlings on UDMH derivatives inducing SOS response. Kinetic curves for the luminescence of *E. coli* MG1655 pColD-lux biosensor cells after the addition of investigated samples: P – water after incubation of *B. inermis* seedlings; P+B – water after incubation of *B. inermis* seedlings together with KK1112 cells; P(UDMH) – UDMH-supplemented water after incubation of *B. inermis* seedlings; B(UDMH) – UDMH-supplemented water after incubation of KK1112 cells; P+B(UDMH) – UDMH-supplemented water after incubation of *B. inermis* seedlings together with KK1112 cells.

In general, results of this measurement are consistent with the results presented in Figure 3C: plant seedlings solely and in consortium with KK1112 bacterial cells efficiently detoxify UDMH-supplemented medium after weeklong incubation. However, one can see that the *E. coli* MG1655 pColD-lux was able to distinguish these two samples. The sample, obtained after incubation of brome seedling with UDMH still possessed DNA damage, which leads to slight rise in MG1655 pColD-lux luminescence. While incubation of brome seedlings together with KK1112 cells in UDMH-supplemented water results in its complete detoxification.

## 4. Discussion

In the work [17], during the development of wastewater biotreatment methods, it was noted that representatives of the genera *Achromobacter* and *Rhodococcus* selected for this purpose could withstand up to 50 mg/L of methylhydrazine. In the work [7], the ability of *Comamonas* sp. isolates to degrade UDMH in soil at concentrations of about 100-600 mg/kg is reported. In our work, a strain of *B. subtilis* resistant to high concentrations of UDMH over 20 g/l was selected with the use of increasing concentrations of UDMH. Judging by the decrease in the alkylating activity of the medium containing 10 g/L UDMH after incubation with *B. subtilis* KK1112 ∼10^9^ cells/mL, we believe that this strain is capable of UDMH biodegradation (Figure 2). It turned out that the reduction of alkylating activity is more effective if a plant-bacteria consortium is used. The greatest efficiency was demonstrated by *B. inermis* seedlings in the presence of *B. subtilis* KK1112 cells (Figures 3 and 4).

With the chosen experimental setup, there may be two scenarios for the effect of plants and *B. subtilis* bacteria on UDMH. First scenario involves accelerating the decomposition of UDMH into safe compounds, which is the ideal outcome. In such case, the activation of the *E. coli* pAlkA-lux biosensor by processed UDMH would not significantly differ from the negative control. Second scenario involves an acceleration of only the first oxidation reaction, i.e. nitrosodimethylamine (NDMA) appears in the medium faster, while the subsequent oxidation occurs spontaneously at a regular rate. In this case, on the contrary, the activation of the biosensor would be observed.

Comparing the data presented in Figures 2 and 3, one can see that activation of biosensor by UDMH-containing solution incubated during a week with *B. subtilis* KK1112 cells at a titer of 5×10^6^ CFU/ml was stronger than activation by samples with higher concentration of UDMH incubated without cells during the same period. This indicates accelerated oxidation of UDMH to NDMA in case of low cell titer. It is the first reaction of UDMH oxidation that can slowly occur without bacteria during oxidation by atmospheric oxygen. Decomposition of NDMA to safe compounds obviously requires an order of magnitude more cells or the presence of plants. The combined effect of bacteria and plants leads to a much more effective reduction in amount of alkylating compounds. This phenomenon is difficult to attribute to anything other than a reduction in the alkylating activity of UDMH derivatives, caused by their more active oxidation to safer compounds. Thus, the presence of *B. subtilis* KK1112 bacteria can promote the biodegradation of high concentrations of UDMH in a shorter time.

It should be noted that the addition of smooth brome seedlings to UDMH results in such a strong drop in the biosensor response that the combined effect of brome and *B. subtilis* KK1112 cells does not result in a further decrease in the response (Figure 3). This can be explained by the fact that after exposure of the UDMH solution to brome seedlings alone (without the addition of bacteria) for a week, NDMA quantity is less than detection limit of biosensor *E. coli* pAlkA-lux. However, some DNA damage properties remain, which could be seen from Figure 5. The absence of this effect in samples, corresponding to the consortium of brome seedlings and KK1112 bacterial cells, is proof of the outstanding effectiveness of this consortium. These results indicate that other DNA damaging substances, in addition to alkylating agents, also do not remain in the medium after incubation of plants in consortium with bacteria. Thus, despite the high efficiency of brome seedlings, the presence of *B. subtilis* KK1112 bacteria could be beneficial for bioremediation of the environment.

Impaired seedling growth is one of the indicators of UDMH exposure. In all studied samples, UDMH exposure caused a significant decrease in the growth of both the root system and the aboveground organs (up to 40% comparing to the control). Even for *B. inermis*, a sharp inhibition of seedling development is observed if *B. subtilis* KK1112 cells are not introduced (Figure 4C), despite the almost complete deactivation of the alkylating potential of UDMH products in the medium (Figure 3C). The addition of the *B. subtilis* KK1112 cells to *B. inermis* seedlings almost completely neutralized the effect of UDMH on seedling development (Figure 4C). The addition of KK1112 to *P. pretense* seedlings had a positive effect on their growth, regardless of the presence of UDMH in the medium (Figures 4B and 4D) and contributed to the detoxification of the UDMH-contaminated medium (Figure 3B). This effect highlights the potential of the KK1112 strain for use in agriculture as a plant probiotic.

In this work, the potential of using selected *B. subtilis* in combination with fodder halophytes, in particular smooth brome, for the detoxification of UDMH-contaminated medium was demonstrated. It could be applied for bioremediation in the arid zones near the Baikonur or other places, where spacecraft launches occurs, like Kourou (Guiana Space Centre), Sriharikota (Satish Dhawan Space Centre), and Jiuquan Satellite Launch Center.

## Author Contributions

Conceptualization, S.Ph.Ch., S.V.B. and I.V.M.; methodology, A.G.K., N.V.K., E.N.B., and I.V.M.; validation, U.S.N., A.G.K., N.V.K., and E.N.B.; formal analysis, S.V.B. and I.V.M.; investigation, U.S.N., A.G.K., N.V.K., and E.N.B.; resources, N.V.K., S.Ph.Ch. and I.V.M.; data curation, S.V.B. and I.V.M.; writing—original draft preparation, N.V.K., E.N.B., S.V.B. and I.V.M.; writing—review and editing, U.S.N., N.V.K., E.N.B., S.V.B. and I.V.M.; visualization, S.V.B.; supervision, I.V.M.; project administration, I.V.M.; funding acquisition, A.G.K., S.V.B., and E.N.B. All authors have read and agreed to the published version of the manuscript.

## Data Availability Statement

All data is available within the manuscript.

## Disclaimer/Publisher’s Note

The statements, opinions and data contained in all publications are solely those of the individual author(s) and contributor(s) and not of Taylor & Francis and/or the editor(s). Taylor & Francis and/or the editor(s) disclaim responsibility for any injury to people or property resulting from any ideas, methods, instructions or products referred to in the content.

