## Supplementary material for "The capability of plant-bacteria consortia to reduce the genotoxicity of unsymmetrical dimethylhydrazine in the environment": Fig S1, Table S1

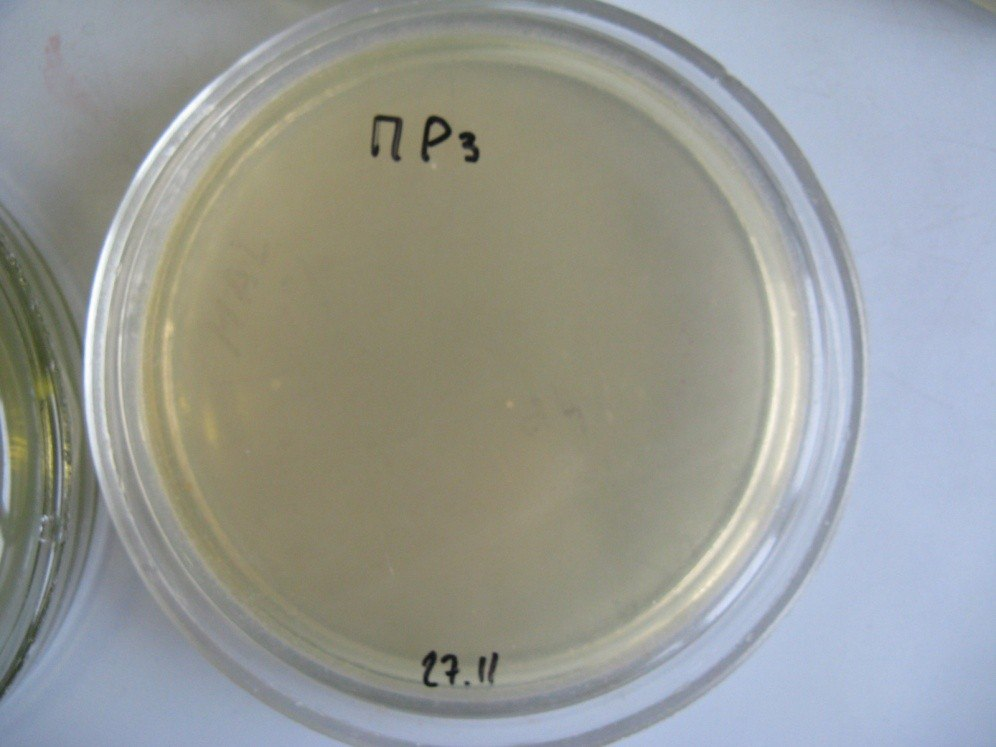

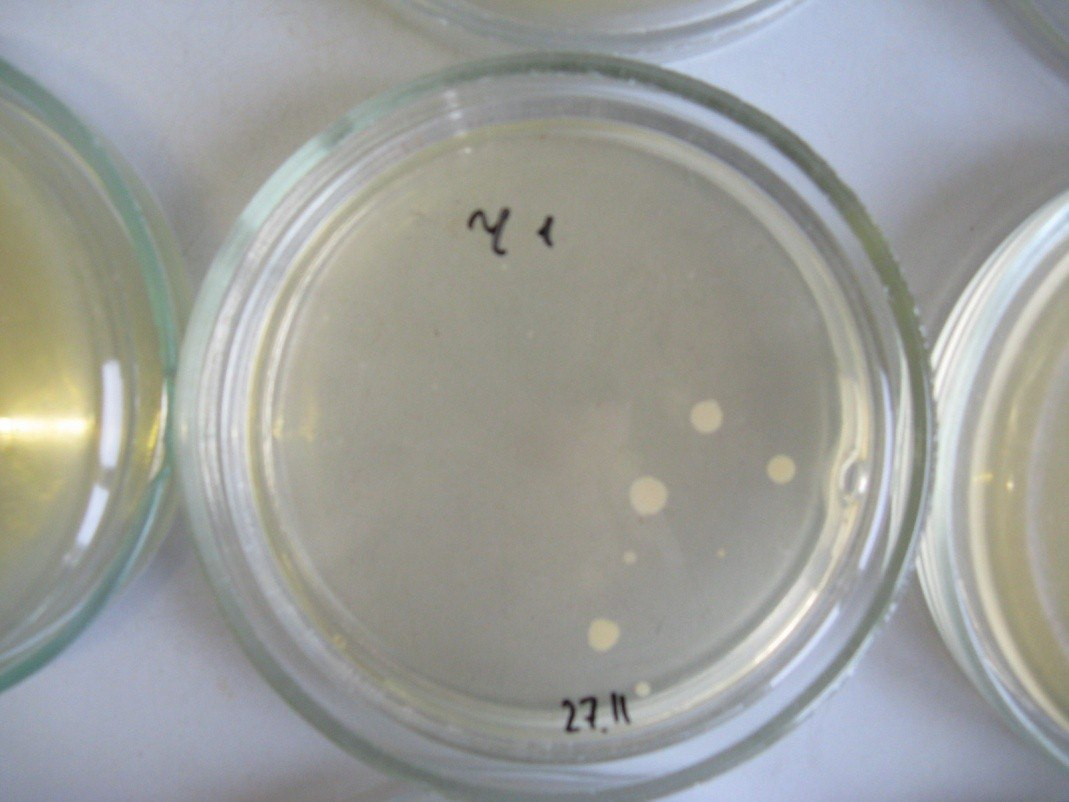

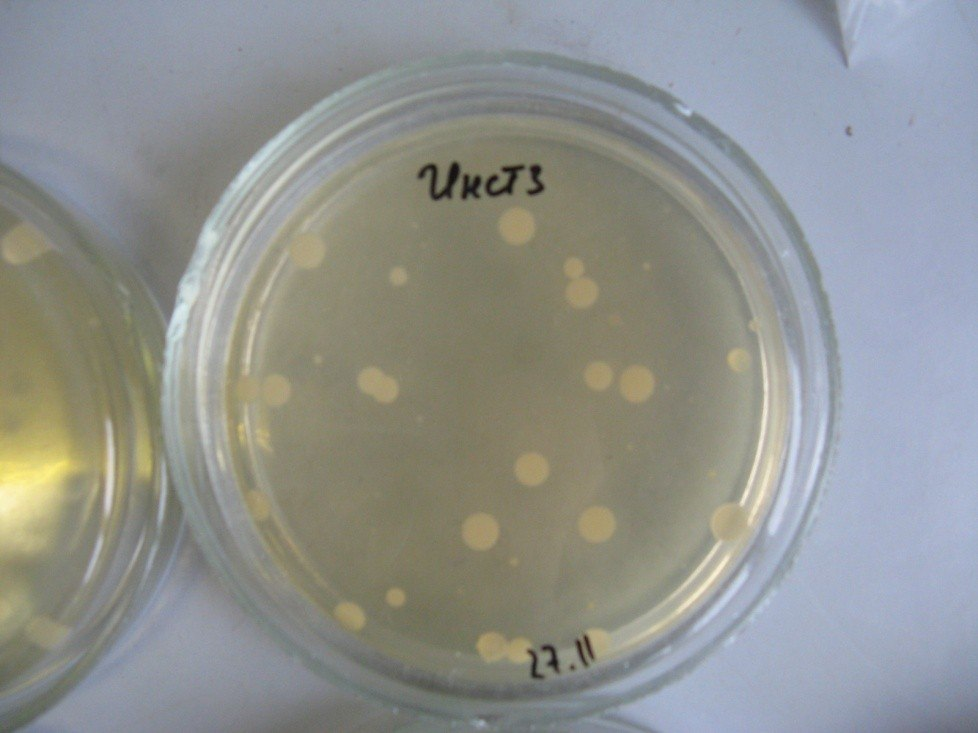

**Figure S1.** The results of plating on LB agar of soil samples, which were diluted in water 1/10 and incubated for a month after supplementation with 10+20 g/L UDMH. From left to right: a sample from the shore of a pond, a chernozem sample, and a sample from the institute's territory.

**Table S1.** Content of viable bacteria in samples after incubation (CFU/mL).

| **Sample** | **Titer, cells/ml** | **Note** |
| --- | --- | --- |
| alfalfa | 1,3×10^4^ | own root microflora |
| timothy-grass | 1,6×10^4^ | own root microflora |
| smooth brome | 5×10^3^ | own root microflora |
| alfalfa + UDMH | ~10^4^ | UDMH additive does not reduce the titer of own microflora |
| timothy-grass + UDMH | 2×10^2^ | UDMH additive reduces the titer of own microflora |
| smooth brome + UDMH | ~10^4^ | UDMH additive does not reduce the titer of own microflora |
| alfalfa + KK1112 | 5×10^7^ | high titer because of *B. subtilis* KK1112 addition |
| timothy-grass + KK1112 | 5×10^6^ | high titer because of *B. subtilis* KK1112 addition |
| smooth brome + KK1112 | 5×10^6^ | high titer because of *B. subtilis* KK1112 addition |
| alfalfa + KK1112 + UDMH | 4×10^7^ | titer does not drop with UDMH addition |
| timothy-grass + KK1112 + UDMH | 6×10^7^ | titer does not drop with UDMH addition |
| smooth brome + KK1112 + UDMH | 8×10^7^ | titer does not drop with UDMH addition |
| KK1112 + UDMH | 5×10^6^ | without plants, KK1112 titer decreases by an order of magnitude |
